# Altered cortical processing of sensory input in Huntington disease mouse models

**DOI:** 10.1101/2021.07.18.452688

**Authors:** Marja D. Sepers, James P. Mackay, Ellen Koch, Dongsheng Xiao, Majid H. Mohajerani, Allan W. Chan, Amy I. Smith-Dijak, Daniel Ramandi, Timothy H. Murphy, Lynn A. Raymond

## Abstract

Huntington disease (HD), a hereditary neurodegenerative disorder, manifests as progressively impaired movement and cognition. Although early abnormalities of neuronal activity in striatum are well established in HD models, there are fewer *in vivo* studies of the cortex. Here, we record local field potentials (LFPs) in YAC128 HD model mice versus wild-type mice. In multiple cortical areas, limb sensory stimulation evokes a greater change in LFP power in YAC128 mice. Mesoscopic imaging using voltage-sensitive dyes reveal more extensive spread of evoked sensory signals across the cortical surface in YAC128 mice. YAC128 layer 2/3 sensory cortical neurons *ex vivo* show increased excitatory events, which could contribute to enhanced sensory responses *in vivo*. Cortical LFP responses to limb stimulation, visual and auditory input are also significantly increased in zQ175 HD mice. Results presented here extend knowledge of HD beyond *ex vivo* studies of individual neurons to the intact cortical network.

## Introduction

Circuit changes and synaptic dysfunction precede neurodegeneration in several adult onset disorders of movement and cognition, including Alzheimer, Parkinson and Huntington disease (HD; reviewed by^1–4^). The most common inherited adult-onset neurodegenerative disorder, HD is a progressive disorder of movement, mood and cognition caused by a CAG triplet repeat expansion greater than 35 in exon 1 of the *HTT* gene, which encodes the protein huntingtin (HTT)^5^. The monogenic inheritance facilitates generation of mouse models with high construct and face validity^6^, and ability to identify gene-expansion carriers in the prodromal stage to enable therapeutic intervention to delay onset of clinical symptoms. Although genetic approaches to lower brain levels of HTT are under investigation in early-stage HD^7^, these may not restore synaptic and circuit function^8^. A better understanding of the mutant HTT (mHTT)-induced mechanisms underlying early alterations in synaptic and circuit function is needed to develop effective treatment for these changes

The earliest neuropathological changes of HD occur in the striatum and in the motor, limbic and associative regions of cortex, which project glutamatergic afferents to the striatum (reviewed by^9,10^), providing input to the basal ganglia-thalamic-cortical loop that selects motor actions and regulates emotional and cognitive behaviours^11^. Abnormalities in cortical-striatal communication that occur prior to neurodegeneration are well documented in mouse models of HD^10–14^, and may, in part, explain early motor incoordination and chorea. Notably, selective knock-down of mHTT in cortical pyramidal neurons ameliorates behavioral phenotypes and improves cortico-striatal synaptic function^15–18^. However, intra-cortical network connectivity and processing in early HD is less well studied.

Although disorders of movement, impairments in learning, and emergence of behaviors associated with depression and anxiety are the subject of numerous studies in HD mice models, less is known about sensory processing. Many patients with HD experience symptoms associated with reduced awareness of their body in space^19^, and posterior cortical regions associated with visuo-spatial processing often show early thinning on MRI^20^. Auditory processing, especially sound source localization and understanding speech in the context of ambient noise, show deficits in HD patients^21^, while older studies suggest sensory perceptual changes and abnormalities of sensory-evoked potentials^22,23^.

To begin to investigate cortical signaling networks in HD, we use sensory stimulation to evoke cortical responses measured on a mesoscale level, using electrophysiological recording from multiple brain regions as well as voltage-sensitive dye imaging *in vivo*, in two different HD mouse models. We find that both YAC128 and zQ175 mice demonstrate enhanced power in sensory responses compared to WT littermates, and *ex vivo* patch clamp recordings suggest increased NMDA receptor-mediated excitatory cortical activity contributes to this enhanced sensory response.

## Results

Cortical sensory processing is not well-studied in HD mouse models; however, changes in cortical regions involved in sensory processing have been reported on structural MRI from prodromal HD gene-expansion carriers^24^. We are interested in mapping cortical network activity on a wide scale, and began by measuring the cortical response to a sensory stimulus because it is time-locked and can be analyzed with fast techniques such as *in vivo* electrophysiology and voltage-sensitive dye imaging to compare genotype responses with precision.

### Response to limb sensory input in YAC128

To evaluate sensory input in YAC128 HD mice, local field potentials (LFP) were recorded in the primary forelimb sensory cortex (FLS1), barrel sensory cortex (BCS1) and motor cortex (M) of 6 month-old mice anesthetized with isoflurane to reduce activity related to voluntary behavior and movement artifacts. Depth of anesthesia was determined by burst suppression due to isoflurane and evaluated for each experiment to ensure equivalence between genotypes (Supplemental Figure 1). Brief subcutaneous electrical stimulation of the contralateral forelimb resulted in an increase in LFP power in FLS1 as shown in representative experiment mean wavelets of stimulation epochs (Figure 1A WT and B YAC128). Instantaneous LFP power in theta (3 to 7 Hz), alpha (7.1 to 12Hz), beta (12.1 to 30Hz) and low gamma (30.1 to 50Hz) frequency bands was calculated by Hilbert transform and normalized to the 2s baseline period before stimulation (Supplemental Figure 2, Figure 1C - J). Genotype summaries of the change in LFP power (area under the curve of the normalized group data for 1.5s following the stimulus) showed a significant increase in YAC128 FLS1 compared to WT (Figure 1; FLS1: frequency p=0.0029, F(3,28) =5.913, genotype p<0.0001, F(1,28)=132.0, no interaction p=0.0512 by 2way ANOVA with Šídák’s multiple comparisons test show in figure). Moreover, the increase in LFP power was not restricted to limb regions of the primary sensory cortex. Strikingly, BCS also show increased change in alpha and gamma LFP power in YAC128 compared to WT (Figure 1 G,H,L, BCS: frequency p<0.0001, F(3,36)=15.71, genotype p<0.0001, F(1,36)=36.65.36 and interaction p=0.3291, F(3,36)=1.185). In the motor cortex, the change power in theta, alpha and beta frequencies were greater in YAC128 (Figure 1I,J,M, frequency p=0.0238, F(3,32)=3.606, genotype p<0.0001, F(1,32)=119.6 and interaction p<0.0001, F(3,32)=10.34 by 2way ANOVA with Šídák’s multiple comparisons test).

**Figure 1.**
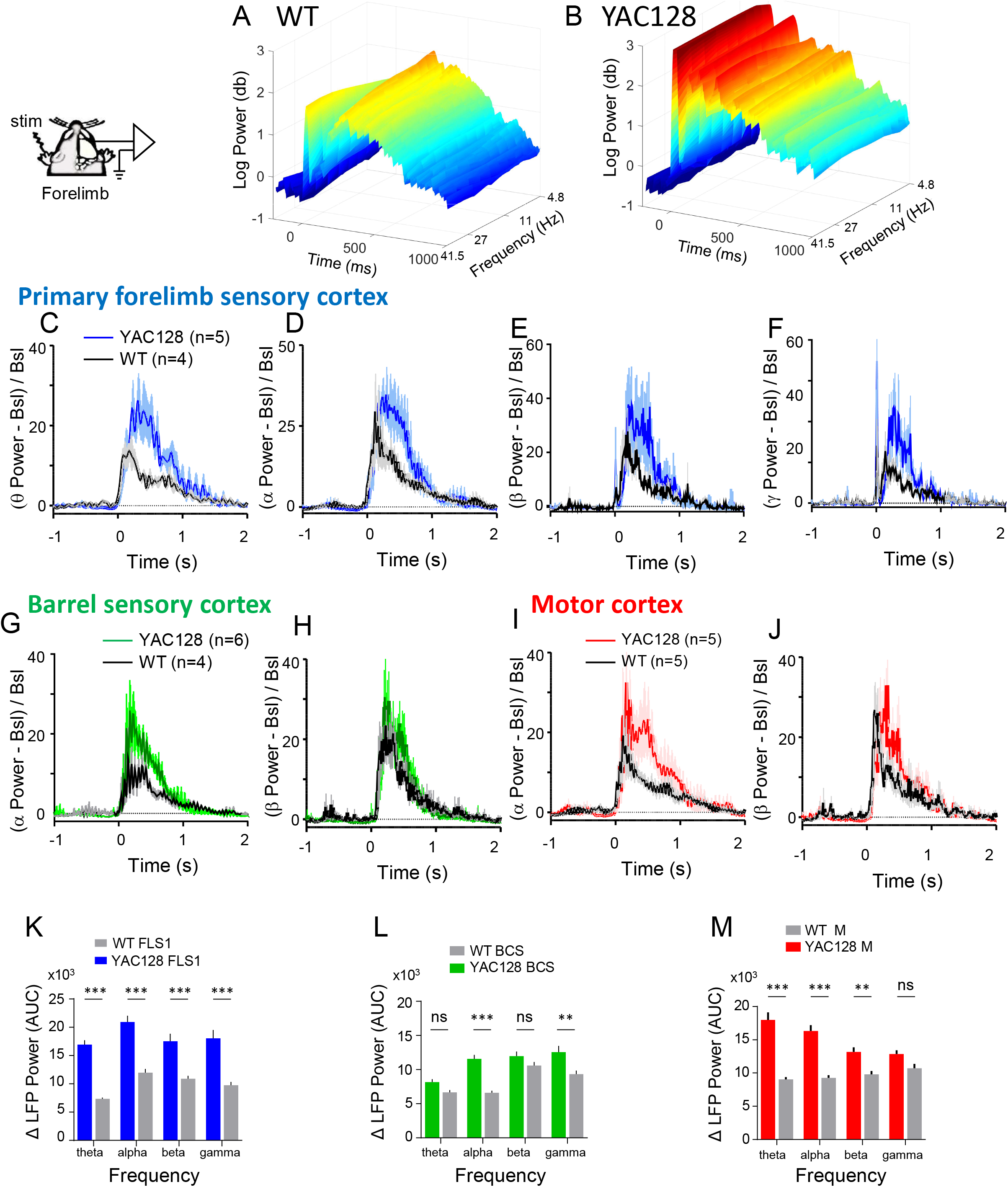
YAC128 HD mice show an augmented response to forelimb stimulation. A and B) Representative local field potential (LFP) wavelets in primary forelimb sensory cortex (FLS1) in a WT and YAC128 mouse with forelimb stimulation at time 0. Time course summary of the change in LFP power (μV^2^) in FLS1 in C) the theta, D) alpha, E) beta and F) gamma frequency bands normalized to baseline. Time course summary of LFP power in Barrel sensory cortex (BCS) at G) alpha and H) beta frequencies and in Motor cortex at I) alpha and J) beta frequencies. Area under the curve (AUC) of change in LFP power in K) FLS1, L) BCS and M) Motor cortex. N = number of mice, results are shown as mean +/− s.e.m. and ***p<0.001, **p<0.01

**Figure 2.**
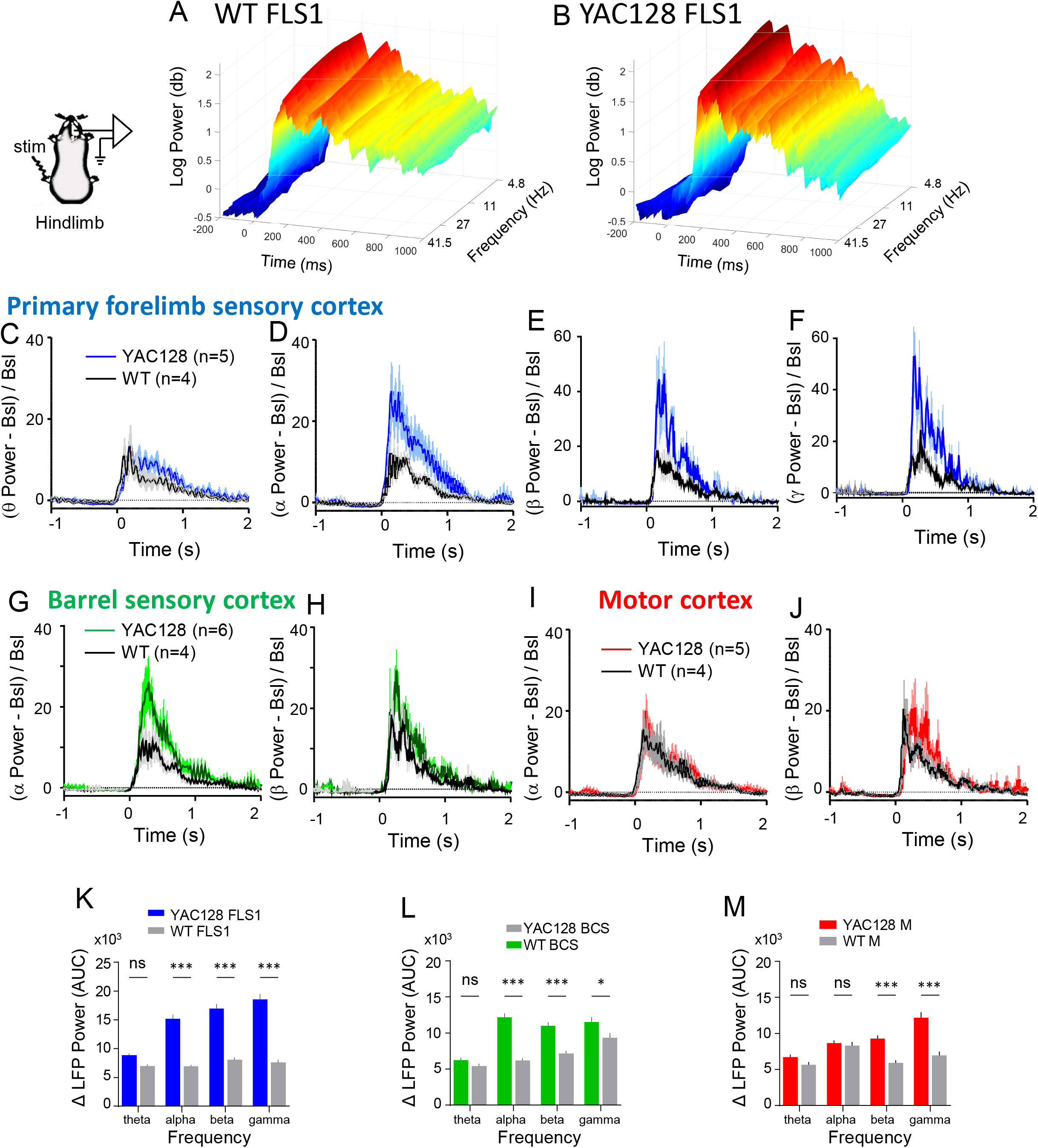
YAC128 HD mice show an augmented response to hindlimb stimulation. A and B. Representative LFP wavelets in FLS1 in a WT and YAC128 mouse with hindlimb stimulation at time 0. Time course summary of the change in LFP power (μV^2^) in FLS1 in C) the theta, D) alpha, E) beta and F) gamma frequency bands normalized to baseline. Time course summary of LFP power in BCS at G) alpha and H) beta frequencies and Motor cortex at I) alpha and J) beta. Area under the curve (AUC) of normalized change in LFP power in K) FLS1, L) BCS and M) Motor cortex. n = number of mice, results are shown as mean +/− s.e.m. and ***p<0.001, *p<0.05

Consistent with a widespread increase in the response to forelimb stimulation, hindlimb stimulation also resulted in a greater change in LFP power in FLS1 at alpha, beta and gamma frequencies (Figure 2 C-F, K; p<0.0001, F(3,28) =27.45 for frequency and p<0.0001, F(1,28)=285.2 for genotype and p<0.0001, F(3,28)=19.3 interaction by 2way ANOVA with Šídák’s multiple comparisons test). BCS and M cortex also show increased alpha (BCS), beta and gamma LFP power over baseline in YAC128 compared to WT (G-J, L, M) following hindlimb stimulation (BCS: frequency p<0.0001, F(3,32) =28.72, genotype p<0.0001, F(1,32)=75.69 and interaction p=0.0002, F(3,32)=9.08; M: frequency p<0.0001, F(3,28)=16.24, genotype p<0.0001, F(1,28)=49.14, interaction p=0.0001, F(3,28)=9.92 by 2way ANOVA).

### VSDI response to limb sensory input in YAC128

To better determine the spatial extent of the cortex activated by limb stimulation, we used mesoscale voltage-sensitive dye (VSD) imaging through a large craniotomy exposing the cortical hemisphere contralateral to the stimulation. As previously described^25^, hindlimb stimulation resulted initially in discrete regional depolarization of primary (HLS1) and secondary (HLS2A and HLS2B) hindlimb sensory cortex in WT with a small expansion to mostly midline cortical areas (Figure 3A). HLS1, FLS1 and secondary sensory areas were functionally determined by the center of activation and used to estimate the position of other cortical areas based on coordinates from the Allen Brain Mouse Reference Atlas. The spread of hindlimb-evoked sensory responses across the cortical surface was markedly more extensive in YAC128 mice compared to WT. In general this manifested as a large non-uniform expansion of the signal into additional areas such as primary BCS along with a longer lasting depolarization (Figure 3A and B). Contralateral hindlimb stimulation in YAC128 mice elicited a transient wave of depolarization encompassing 21.87 ± 2.17 mm^2^ (n=5) of the cortical surface (defined as pixels with a ΔF/F response at least 5x baseline RMS noise) compared to a significantly smaller 6.85 ± 2.61 mm^2^ (n=4) response in WT mice (p=0.0029, t=4.473, df=7 by 2-tailed unpaired t-test).

**Figure 3.**
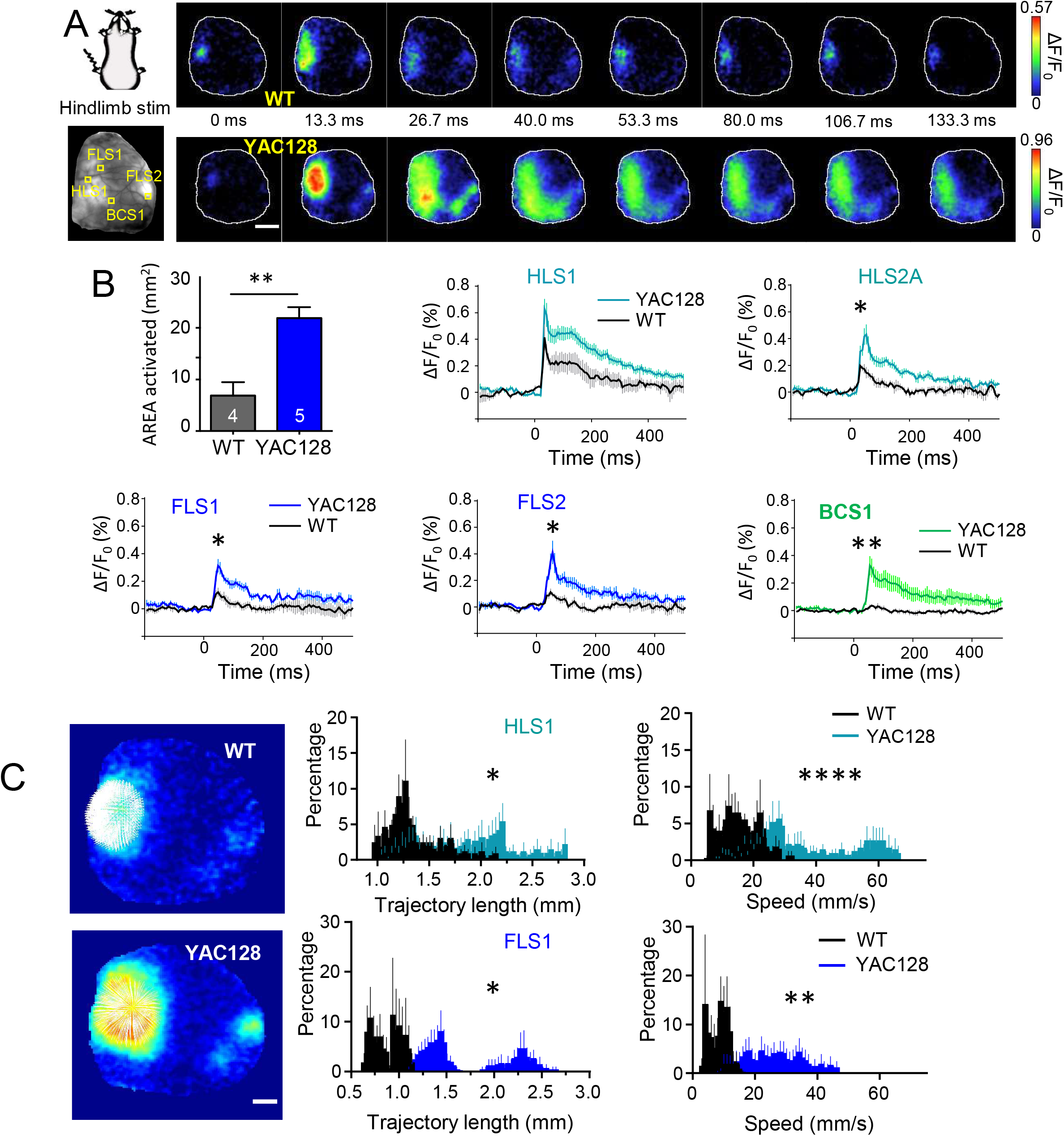
Voltage sensitive dye imaging of the cortex with limb stimulation reveals increase in the area activated, trajectory and speed of the spread of the response in YAC128 mice. A) Time series of cortical wide-field voltage-sensitive dye imaging (VSDI) signals (ΔF/F), from the right hemisphere of representative wildtype (top) and YAC128 (bottom) mice, during contralateral hindlimb (HL) sensory-stimulation scale:2mm. Note evoked signals, reflecting neuronal depolarization, propagate more extensively across the YAC128 cortical surface. B) Cortical surface area activated following hindlimb stimulation in WT (grey) (n=4) and YAC128 (blue) (n=5) mice and time course of activation in cortical areas following stimulation at time 0ms. Significant differences between genotypes of the peak response are indicated. C) Example images of VSD responses originating from primary sensory areas in response to hindlimb stimulation (left, scale:1mm). Pixel trajectory length and maximum speed in response to hindlimb stimulation were both increased in YAC128 compared to WT for hindlimb primary sensory area (top panels) and forelimb primary sensory area (bottom panels). *p<0.05, **p<0.01, ****p<0.0001

The spatiotemporal spread of VSD signals in mesoscale imaging makes these data amenable to optical flow analysis^26^. This approach calculates velocity vector fields to quantify the speed and direction of motion^27^. In the context of our data, we can measure the trajectory, direction and speed of the spread of neural activity across the cortex, represented by changes in the brightness of pixels over time and space. We quantified VSD cortical dynamics with the Optical Flow Analysis Toolbox (OFAMM)^27^ (available at http://lethbridgebraindynamics.com/ofamm/) which revealed an increase in trajectory length (HLS1: p=0.0247, K-S D=0.2198; FLS1: p=0.0238, K-S D=0.1773) and temporal speed (HLS1: p<0.0001, K-S D=0.4648; FLS1: p-0.0041, K-S D=0.2475) in YAC128 from both HLS1 and FLS1 in response to hindlimb stimulation (Figure 3C). Secondary HL and FL areas, as well as primary and secondary barrel cortex also showed increased trajectory length (HLS2A: p=0.0075, K-S D=0.2149; HLS2B: p<0.0001, K-S D=0.3967; FLS2: p=0.0075, K-S D=0.0075; BCS1: p<0.0001, K-S D=0.2893; BCS2: p-0.0031, K-S D=0.2314) and maximum temporal speed (HLS2A: p<0.0001, K-S D=0.7750; HLS2B: p<0.0001, K-S D=0.5614; FLS2: p=0.0561 (ns), K-S D=0.1881; BCS1: p=0.0054, K-S D=0.3115; BCS2: p<0.0001, K-S D=0.6087) in YAC128 after hindlimb stimulation (Supplemental Figure 3).

In contrast to hindlimb stimulation, responses to forelimb stimulation resulted in widespread cortical depolarization that did not significantly differ between genotypes: (18.78 ± 5.35 mm^2^ (n=5) and 21.68 ± 1.66 mm^2^ (n=4) in YAC128 and WT mice respectively; p=0.6554, t=0.4660, df=7 by 2-tailed unpaired t-test). However, HLS1, FLS1 and BCS1 depolarization in YAC128 all showed higher correlation with other cortical areas during the 500ms following hindlimb stimulation compared to WT as shown by seed-pixel correlation maps and regional correlation matrices (Supplemental Figure 4A). In addition, correlation between cortical areas following whisker stimulation was abnormally high in YAC128 compared to WT, suggesting that other types of sensory stimulation are augmented in YAC128 mice (Supplemental Figure 4C).

### Synaptic events *ex vivo* in YAC128

In HD models, the cortex and striatum have an imbalance in excitation and inhibition with increased glutamate signaling at extrasynaptic NMDA-type glutamate receptors (NMDAR;^4,13^). Furthermore, HD stage-dependent changes in inhibitory and excitatory input to cortical pyramidal neurons have been reported in acute brain slice recordings from R6/2 and YAC128 HD mice^28,29^. The augmented sensory response in 6 month-old YAC128 mice shown here could result from increased excitation or decreased inhibition in cortical circuits. To investigate these possibilities in acute brain slices, we conducted whole-cell voltage clamp experiments to measure excitatory and inhibitory synaptic responses in the sensory cortex of 6 month-old YAC128 mice. The cell capacitance and membrane resistance of layer 2/3 pyramidal neurons was similar in WT (n=24(7), capacitance 102 +/− 8.07 pF; resistance 146.1 +/− 18.14 MΩ) and YAC128 (n=19(6), capacitance 101.6 +/− 11.13 pF, NS p=0.97, t=0.03, df=41 by unpaired t-test; resistance 188.5 +/− 31.08 MΩ; NS p=0.23, t=1.23, df=41). Excitatory postsynaptic currents (EPSCs) in layer 2/3 pyramidal neurons were evoked by a short train of 10 stimulations at 20Hz with a microelectrode placed 300μm ventral. NMDAR-mediated EPSC were isolated by holding the cells at +30mV while blocking AMPAR with CNQX and GABA_A_R with PTX. There was no difference between genotypes in the amplitude of EPSCs evoked by 20Hz train stimulation (Supplemental Figure 5, p = 0.15 unpaired t test, t=1.11, df=9), but on some trials electrogenic events occurred that were > 5x larger in amplitude and obscured the baseline EPSC (Figure 4A). In order to compare evoked EPSCs without large amplitude events, the lowest stimulation intensity required to evoke synaptic responses was used (not different between genotypes). However, more YAC128 neurons than WT neurons showed these large amplitude events that have previously been described as “NMDAR spikes”^30–32^ (*p=0.04, Chi-square=4.295, df=1, n=10(6) YAC128 and n=11(7) WT) following minimal stimulation and they also occurred spontaneously (Figure 4 B). When glutamate uptake was blocked by dl-TBOA (Figure 4D and E), YAC128 neurons showed a greater increase in event amplitude (**p=0.003 Kruskal-Wallis ANOVA; n= 7(6) WT and n= 8(6) YAC128; with Dunn’s multiple comparisons in YAC128 control vs. TBOA *p=0.03) and frequency (Kruskal-Wallis ANOVA, approximate **p=0.004 with Dunn’s multiple comparisons in YAC128 *p=0.04,) and compared to WT neurons (Kruskal-Wallis ANOVA as above, Dunn’s multiple comparison before vs after TBOA in WT p=0.14 for frequency and p=0.08 for amplitude).

**Figure 4.**
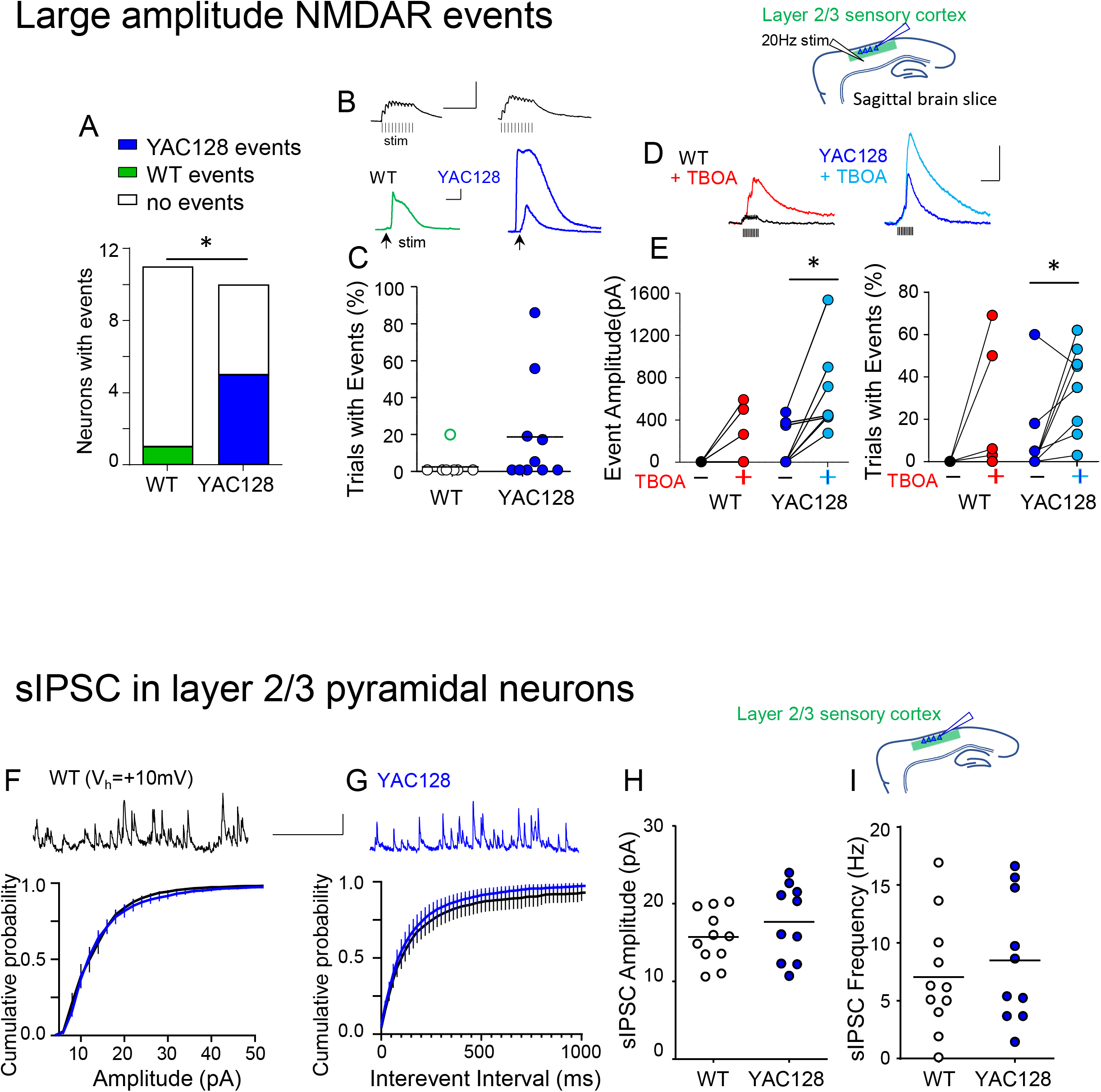
Neurons in the sensory cortex of YAC128 mice show more evoked large excitatory events than WT but similar spontaneous inhibitory post synaptic currents (sIPSC). (top) diagram of electrode positions in acute brain slice. A) Compared to WT, more YAC128 neurons expressed NMDAR events with amplitude greater than 200pA. B) representative synaptic responses to 20Hz stimulation (above) and NMDAR events evoked by stimulation (below) in WT (green) and YAC128 (blue) Note difference in scalebar showing 200pA and 0.5s. C) Frequency of NMDAR events for each neuron. D) representative responses to stimulation and E) summary of event amplitude and frequency before and after application of TBOA to induce glutamate spillover. F-I) Representative IPSCs traces with scalebar: 20pA,1s. and Cumulative probability of F) amplitude and G) frequency shown as mean +/− s.e.m. H) sIPSC amplitudes and I) frequency showing no difference between WT and YAC128 neurons from layer 2/3 (circles represent individual cells with line at the group mean). *p<0.05

In contrast to the difference in excitatory events seen in YAC128 layer 2/3 pyramidal neurons, spontaneous inhibitory postsynaptic currents (sIPSCs) frequency and amplitude were similar in these neurons from both WT and YAC128 mice recorded at +10mV in the sensory cortex (Figure 4B). We also recorded miniature IPSCs in layer 5 pyramidal cells and found no significant difference between WT and YAC128 in either the mean frequency (WT 14.86Hz +/− 1.28 vs 13.73Hz +/− 1.68 YAC128, p=0.59, t=0.54, df=22, by unpaired two tailed t-test with n=13(5) and n = 11(5) (respectively) or amplitude (WT 54.13 pA +/− 5.34 vs YAC128 43.67pA +/− 7.34, p=0.25, t=1.176, df=22). Together, these data suggest an increase in excitatory input rather than altered inhibition to cortical layer 2/3 pyramidal neurons contributes to enhanced cortical spread of sensory responses in YAC128 mice.

### Limb sensory input in zQ175

To determine if the altered sensory response observed in YAC128 mice was found in other HD mouse models, we measured LFP power in zQ175 HD mice. The level of anesthesia for these experiments was lower than that used in YAC128 mice (Supplemental Figure1) to facilitate comparison of multi-unit (MU) activity between groups. Consistent with the results found in YAC128 mice, zQ175 LFP power responses to both hindlimb and forelimb stimulation were augmented compared to WT littermates in FLS1 (Figure 5A-D,J,K,hindlimb: frequency p=0.0042, F(3,36)=5.239, genotype p<0.0001, F(1,36)=119.1. Forelimb: frequency p=0.0002, F(3,36)=8.588, genotype p<0.0001, F(1,36)=123.5 by 2way ANOVA). The LFP response was also augmented in zQ175 BCS and M1 to hindlimb stimulation (Figure 5F and H. BCS: frequency p=0.0673, F(3,36)=2.598 and genotype p<0.0001, F(1,36)=49.46. M1: frequency NS p=0.1192, F(3,40)=2.072 and genotype p<0.0001, F(1,40)=43.72 with interaction p=0.0334, F(3,40)=3.201) and forelimb stimulation (Figure 5M and O; BCS: frequency p=0.0025, F(3,32)=5.909 and genotype p<0.0001, F(1,32)=78.67 with interaction p=0.0006, F(3,32)=7.554. M1: frequency p<0.0001, F(3,32)=9.809 and genotype p<0.0001, F(1,32)=122.4 and interaction p=0.0001, F(3,32)=9.543). Interestingly, although the probability of MU activity increased in FLS1, M1 and BCS following hindlimb stimulation (Figure 5E,G,I, Supplementary Figure 6), there was no difference between genotypes (FLS1: time p=0.0111, F(3.55, 17.78)=4.674 and genotype p=0.6709, F(1,5)=0.2034. BCS: time p<0.0001, F(49,343)=7.717 and genotype p=0.0539, F(1,7)=5.351. M1: time p<0.0001, F(49,343)=7.843 and genotype p=0.2929, F(1,7)=1.293 by 2way RM ANOVA).

**Figure 5.**
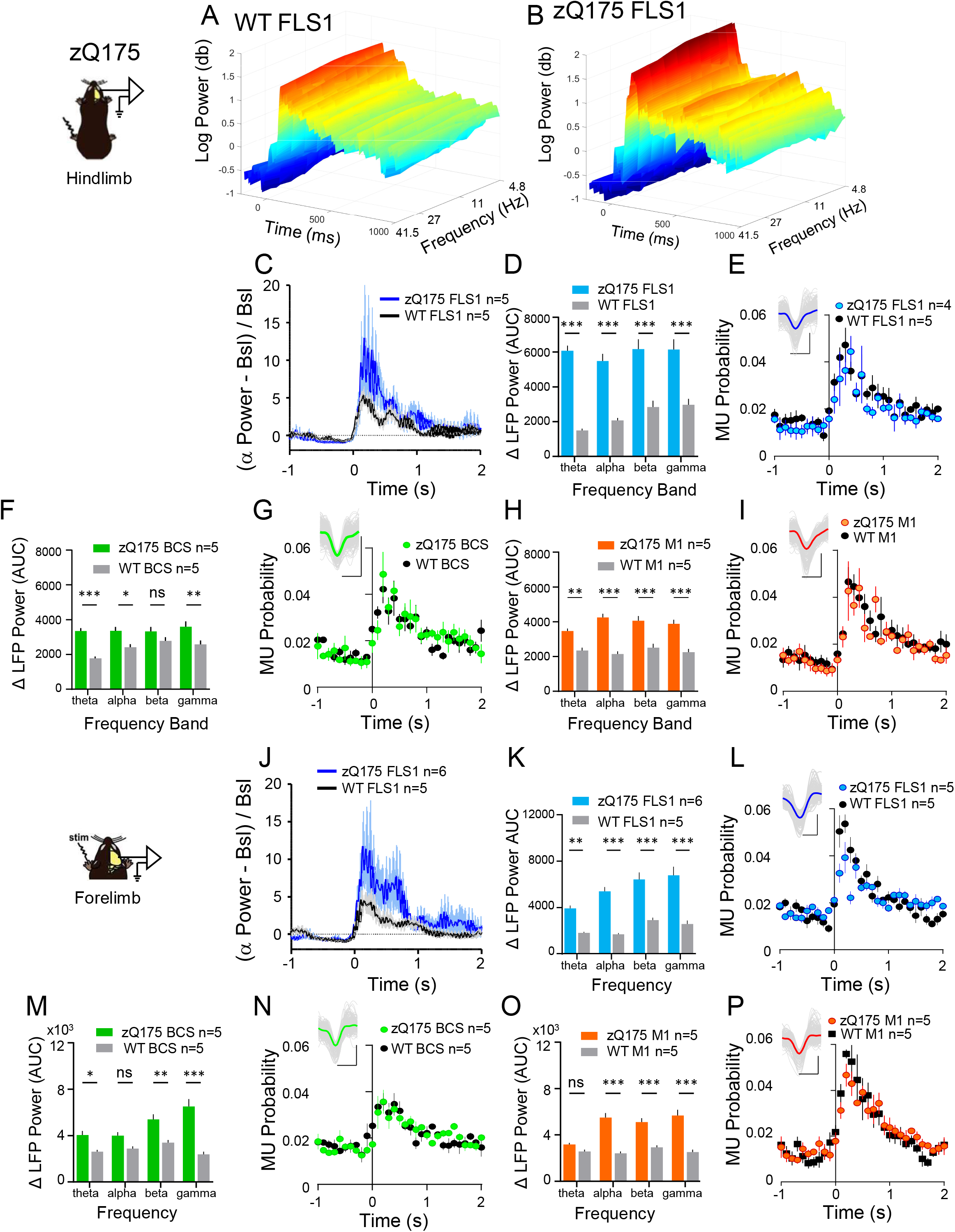
zQ175 HD mice under anesthetic show an augmented LFP power with limb stimulation but multi-unit neuronal activity is similar to WT cortical areas. A and B) representative wavelets from FLS1 in a WT and zQ175 mouse with hindlimb stimulation at time 0. C) Time course summary of FLS1 normalized alpha power in WT (black) and zQ175 (blue). D) FLS1 change in LFP power in frequency bands shown as area from 0 to 1.5s following stimulation as shown for alpha in C. E) Local neuronal activity shown as probability of Multi-unit (MU) events in FLS1. Inset shows representative events and mean waveform for one experiment. F and G) BCS LFP power summary and MU probability in response to hindlimb stimulation. H and I) M1 LFP power summary and MU probability in response to hindlimb stimulation. J-P) LFP power and MU probability as above in response to forelimb stimulation. Data shown as mean +/− s.e.m. *p<0.05

### Auditory and Visual sensory input in zQ175

Unlike YAC128 mice that are on a visually impaired FVB/N background^33^, zQ175 mice on a C57/Bl6 background could be tested with a visual stimulus. As shown in Figure 6, a 10ms blue LED flash evoked a greater response in zQ175 than WT in FLS1, M1 and BCS1 (FLS1: frequency p=0.3785, F(3,32)=1.063 and genotype p<0.0001, F(1,32)=180.0. BCS1: frequency p=0.1217, F(3,24)=2.139 and genotype p<0.0001, F(1,24)=79.75. M1: frequency p=0.4351, F(3,28)=0.9387 and genotype p<0.0001, F(1,28)=119.1). The LFP power increase and MU activity responses to visual stimulation appeared smaller and slower than other sensory responses. A brief auditory stimulus also resulted in a significantly increased LFP power in zQ175 over WT in all areas tested (Figure 6H,I,K,M. FLS1: frequency p=0.0003, F(3,36)=8.209 and genotype p<0.0001, F(1,36)=98.73 with interaction p=0.0065, F(3,36)=4.805. BCS1: frequency p=0.0008, F(3,32)=7.212 and genotype p<0.0001, F(1,32)=141.9 and interaction p=0.0196, F(3,32)=3.794. M1: frequency p<0.0001, F(3,32)=11.26 and genotype p<0.0001, F(1,32)=77.37 and interaction p=0.0774, F(3,32)=2.498). MU activity increased following both visual and auditory stimulation (auditory FLS1 time p<0.0001, F(49,343)=7.356; BCS time p<0.0001, F(49,343)=4.936; M1 time p<0.0001, F(49,343)=7.270) and unlike limb stimulation there was significant interaction between time and genotype (FLS1 genotype p=0.1491, F(1,7)=2.627 with interaction p=0.0062, F(49,343)=1.645. BCS genotype p=0.5024, F(1,7)=0.5 and interaction p<0.0001, F(49,343)=2.151. M1 genotype p=0.2098, F(1,7)=1.906 with interaction p=0.0280, F(49,343)=1.467) suggesting a more varied or complex local response.

**Figure 6.**
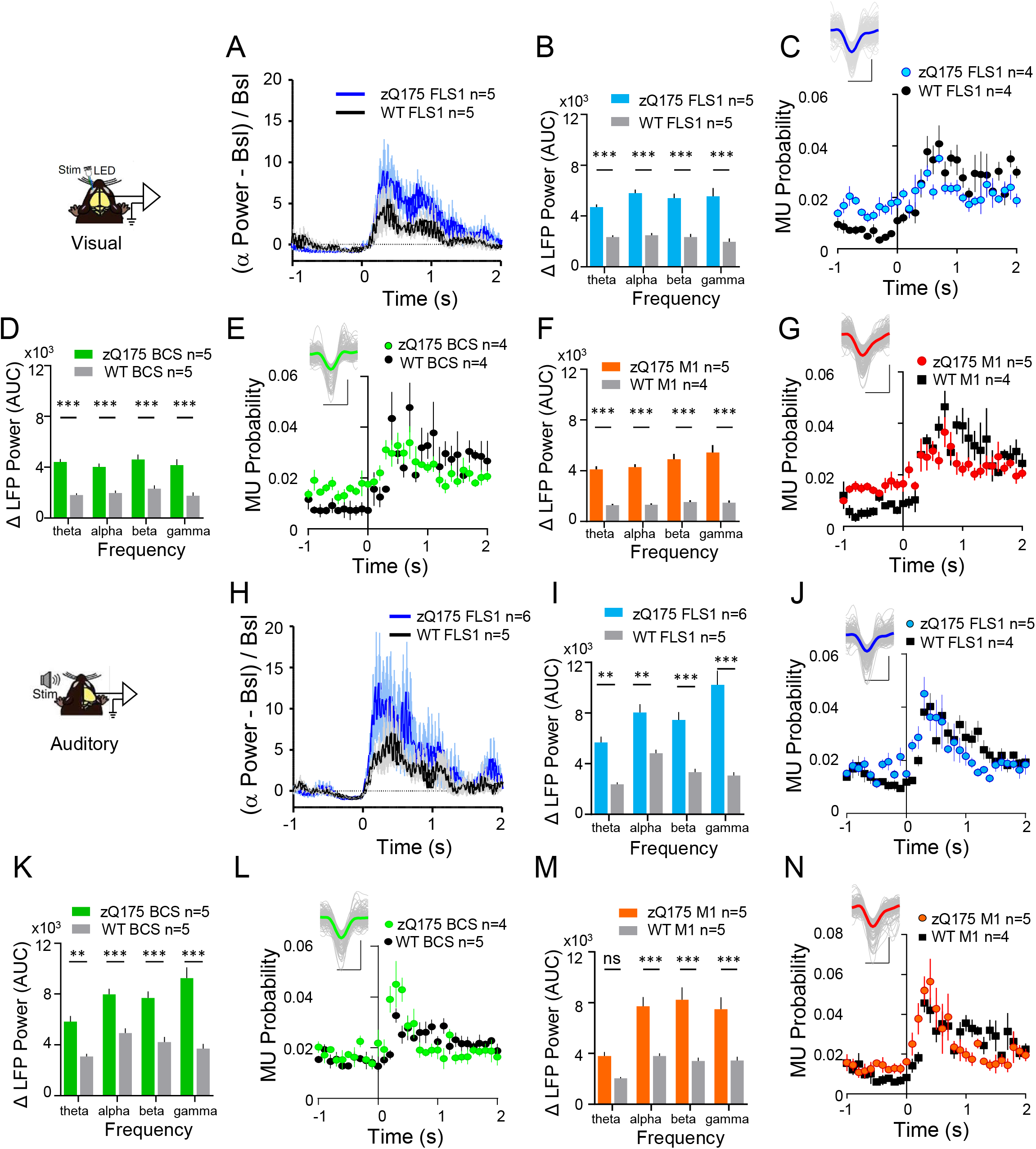
zQ175 HD mice show an augmented LFP power in response to auditory and visual sensory input. A) Time course summary of LFP power at alpha frequencies in WT (black) and zQ175 (blue) FLS1 with a 10ms blue led flash at time 0. B) Comparison of LFP power in FLS1 WT and zQ175 showing greater activation in all frequency bands. C) MU event probability increases after visual stimulation but shows no significant difference between genotypes. D and E) LFP power increase in response to visual stimulation and MU event probability in BCS. F and G) LFP power increase and MU event probability in M1. H-N) zQ175 and WT LFP response to auditory sensory stimulation shown as above. Data shown as mean +/− s.e.m. **p<0.01,***p<0.001

Overall, the significantly increased sensory response in LFP power in zQ175 is not reflected in MU activity suggesting that local neuronal post-synaptic supra-threshold responses are not the primary driver of widespread cortical activation. Indeed, the similarity of responses in LFP power and VSD imaging correlation between cortical areas is consistent with increased connectivity and a more global sensory response in both YAC128 and zQ175 mice.

## Discussion

Previously published studies suggest altered sensory processing in patients with Huntington disease^19,21–23^, but these changes and their underlying mechanisms have not been fully investigated. Aberrant cortical processing of sensory input could impact accuracy of movement and also impair cognition, key areas of clinical decline in patients with HD. Here we report that cortical responses to sensory stimulation, as measured by *in vivo* brain imaging or electrophysiological approaches, are augmented in two HD mouse models, YAC128 and zQ175, compared to WT littermates. Multiple cortical areas were depolarized in YAC128 in contrast to the discrete spatial response in WT mice shown here by VSDI. Results for hindlimb stimulation as observed by VSDI and LFP recordings were remarkably complementary, with both methods showing augmentation of the sensory response by cortical dynamics in the HD model mice.

The spread of sensory-evoked signals across the cortical surface (sensory-spread), measured with VSDI or other methods, has been documented in numerous mammalian species ranging from rodents to primates^34–36^. This phenomenon involves the coordinated activity of thousands of cortical neurons and is thought to be subserved by a diffuse network of inter-cortical projections, which extend radially from individual cortical columns in all directions^37,38^. The physiological role(s) of sensory-spread are incompletely understood^39–41^. However, the widespread signals across the cortical surface, often irrespective of function boundaries, suggests diverse roles in cortical integration. Although we have examined cortical activity exclusively in anaesthetized animals, sensory spread is observed in awake animals^26,42^.

In WT animals the extent of the sensory spread (measured with VSDI) varied considerably with the modality tested. In contrast, YAC128 mice show consistently large areas activated with the maximal hindlimb sensory-spread more than triple that of WT. Consistent with this, coherence between cortical areas was greater in YAC128 than WT mice not only following hindlimb stimulation, but also whisker stimulation. Similarly, both YAC128 and zQ175 mice had greater responses to multiple sensory modalities compared to WT in motor and sensory cortical areas, including forelimb stimulation when responses were measured by recording local field potentials, which are more sensitive to high frequency oscillations.

At the level of cortical synapses, impaired balance of inhibition and excitation could contribute to the increased sensory spread in YAC128. Sensory cortical spontaneous IPSCs are reduced in older symptomatic R6/2 mice and EPSCs are more frequent^28,43^. However, the same study showed an increase in IPSC frequency in the CAG140 and YAC128 models. Previous studies of neurons in the motor cortex showed a decrease in IPSC amplitude with an increase in frequency in 12 month-old zQ175 mice^29^. A decrease in inhibition in layer 2/3 of the motor cortex is also shown by *in vivo* calcium imaging and *ex vivo* immunohistochemistry in HD mouse models and patients^44,45^. These studies have shown conflicting results as to how inhibition is affected in the cortex in HD, and likely differ depending on the region studied. Here, in HD mice that exhibit aberrant sensory stimulus-induced activation of both the sensory and the motor cortex *in vivo*, our *ex vivo* experiments show no difference between WT and YAC128 mice in the amplitude or frequency of sIPSCs recorded from sensory cortex pyramidal neurons in layer 2/3 or of mIPSCs in layer 5. Although YAC128 mice at 6 months of age do show an HD-like phenotype and synaptic deficits, it is possible that a change in IPSCs occurs in older YAC128 mice.

Interestingly, cortical neurons in R6/2 mice show enhanced spontaneous, large amplitude “complex” events that increase in frequency and duration compared to WT as the mice age^28,29^. Here we report that YAC128 neurons display larger amplitude events in response to stimulation of excitatory afferents than WT. Although the experimental conditions were different in the R6/2 study, it is striking that both HD models exhibit unlooked-for large amplitude excitatory events. We and others have previously shown increased excitatory transmission and extrasynaptic NMDAR function in the striatum in HD models^12,13,46–48^. The large amplitude events shown here are consistent with augmented extrasynaptic NMDAR-mediated events in the cortex of YAC128, since they increase with glutamate spillover and occur in the presence of AMPA and GABA_A_ receptor blockers. It is possible that these events occur due to network bursting and synaptic integration^30^ or by astrocytic release of glutamate following stimulation^49^. Astrocytes also contribute to hyperexcitability in the striatum in HD models, although fewer studies in HD models look at astrocytes in the cortex^50^. Future studies will investigate the mechanism of these events and their relation to sensory responses *in vivo*.

In addition to the increased spread of sensory-evoked VSD responses in YAC128 cortex, the optical flow analysis revealed a greater maximum temporal speed of signal propagation. Previous work has shown that sensory cortical neurons of R6/1 and R6/2 mice have increased input resistance, decreased cell capacitance, and a depolarized membrane potential at symptomatic ages^28,51^. Cortical pyramidal neurons from YAC128 and CAG140 mice also exhibit increased input resistance starting at 6 or 12 months of age, but normal resting membrane potential and cell capacitance^28^. Although those previously published data suggest that changes to membrane properties of cortical neurons in the HD brain could explain the observed increase in propagation speed, our data show no significant difference in membrane capacitance or resistance in layer 2/3 cortical pyramidal neurons from 6 month-old YAC128 vs. WT mice.

Subthreshold signals in dendritic processes, which represent the majority of the neuronal surface area, appear to predominantly mediate the sensory-spread^34,52^. Based on this, it’s perhaps not surprising that the enhanced sensory-spread in zQ175 mice was not associated with increased cortical multi-unit firing probability. Regardless of the underlying mechanisms, it seems that the aberrant cortical sensory processing is more dependent on alterations to network synchrony than single-neuron bursting activity. However, recordings with higher density electrodes would be required to further examine the sensory-evoked activity of individual cortical neurons in relation to their relative anatomical positions. It is noteworthy to mention that we noticed a region- and stimulus-dependent variability in the spiking activity between animals that could result from the relative locations of cortical point spreads to the recording sites^53^.

Given that a correlated noise over large cortical areas can decrease stimulus acuity^54^, the augmented cortical sensory response in HD mice could be detrimental to performance of sensory-motor tasks and contribute to impaired motor learning. In fact, inhibition in the sensory cortex is important for hand grasping in humans^55^, and HD patients typically show deficits in reaching and grasping movements^56^. Human EEG studies show that suppression of gamma power in the visual cortex modulates reactions to sensory input^57^, suggesting that increased gamma power in other cortical areas could also impair reactions to sensory input. It is interesting to note that in YAC128 mice, hindlimb sensory responses of higher frequency LFP in the beta and gamma range were more augmented than theta frequency power. Similarly, aberrantly increased gamma oscillations in awake behaving mice have been shown in the cortex and striatum of the R6/2 mouse model of HD^43^.

Taken together with the increased LFP response to visual and auditory stimulation in zQ175, our results showing enhanced VSDI and LFP response to limb stimulation demonstrate consistently aberrant cortical dynamics that generalize across HD models and sensory modalities. These results in conjunction with our *ex vivo* slice data suggest subthreshold synaptic depolarizations, perhaps mediated by increased extrasynaptic NMDA receptor expression, underlie the increased sensory-spread in HD mice.

## Methods

### Animals

All procedures were performed in accordance with the Canadian Council on Animal Care and University of British Columbia Animal Care Committee regulations (approved under protocol A19-0076). Mice were group housed with 2 to 4 mice per cage on a 12h light, 12h dark cycle. Water and standard laboratory mouse diet were available ad libitum. Male 5 to 6 month old YAC128 Line 53 and their wild-type FVB littermates^58^, and zQ175DN mice (https://www.jax.org/strain/029928;^59^) and their wild-type C57/B6 littermates were implanted with electrodes as below and allowed to recover for 1 to 4 weeks before experiments.

### Electrode Implant surgery

Mice were anesthetized with isoflurane at 3% for induction then reduced to 1.5 to 2 % for stereotaxic surgery. The eyes were covered with eye lubricant (Lacrilube; www.well.ca) and body temperature was maintained at 37⁰C using a heating pad with a feedback thermistor. A skin flap extending over the dorsal cortex was cut and removed. Fascia or connective tissue was lightly scraped away from the skull and small (< 1 mm diameter) holes were drilled through the skull, using a high-speed dental drill with sterile bit, over the cortex. Twisted tungsten wire tetrodes (25μm diameter, California Fine Wire Co.) -- typically 3 -- were directed toward the center of burr holes and placed in the cortex (depth: 500 μm) using a motorized micromanipulator (MP-225, Sutter Instrument Co.). Electrodes were implanted in the cortex at the following coordinates in mm relative to Bregma; primary forelimb sensory cortex (FLS1: AP 0.5, ML 2.25, DV 0.5), barrel sensory cortex (BCS1: AP 0.1, ML 3, DV 1) and motor cortex (M1: AP 1, ML 1.5, DV 0.5).

Miniature connectors (2 × 2 × 2 mm) were cemented to the skull (with dental adhesive). Ground and reference electrodes (silver wire) were fixed onto the surface of the posterior skull. Prior to implantation, tetrodes were painted with fluorescent 1,1-dioctadecyl-3,3,3,3-tetramethylindocarbocyanine perchlorate (DiI, ~10% in dimethylfuran, Molecular Probes, Eugene, OR) and the solvent allowed to evaporate. For histology, immediately following the experiment, animals were decapitated and the brain fixed in 4% paraformaldehyde. The brains were sliced on a vibratome and diI labeling counter-stained with DAPI was used to identify the tetrode tract and confirm the approximate cortical location.

### *In vivo* electrophysiology

Mice were anesthetized with 1 - 2% Isoflurane and body temperature maintained at 37⁰C with heating pad and thermistor (Harvard Apparatus). To stimulate forelimb and hindlimb, a 1ms pulse of 0.5 - 1mA was delivered by thin acupuncture needles (0.14mm) inserted subcutaneously into the paw. In zQ175 experiments visual stimulation was a 10ms flash of blue light and auditory stimulation was a 1ms broad frequency spectrum ‘chirp’ produced by a TTL pulse to a Piezo Element (Adafruit). Responses were calculated from 10 - 20 trials of stimulation, each trial separated by 10s. LFP signal was recorded at 25kHz and filtered (0.1 to 1000Hz) using a 16-channel data acquisition system with a 1200 gain (USB-ME16-FAI-System, Multi Channel Systems or an RHD2132 amplifier chip and Intan recording controller). LFP power analysis of one tetrode wire from each area was performed in MATLAB (2019; Mathworks, Natick, MA). Isoflurane burst suppression was evaluated to ensure groups were at similar anesthetic levels by measuring the mean duration between bursts (Supplemental Figure 1). If the level of anesthetic changed during the experiment, those stimulation trials were excluded from the analysis. The Morlet wavelets (6 cycle) of each epoch were averaged for visualization of the experiment. LFP recordings were band-pass filtered^60^ for theta (3 to 7Hz), alpha (7.1 to 12Hz), beta (12.1 to 30Hz) and low gamma (30.1 to 50Hz) frequencies followed by isolating epochs of 2s before and 3s after stimulation with no bursting activity in the 200ms before stimulation (automatically determined by an absolute value greater than 200μV in the unfiltered data). 10 to 20 trials were analysed for each experiment after exclusion criteria were applied as above. Hilbert transform for instantaneous power calculated (using the MATLAB function) for each individual trial epoch and the results averaged as shown in Supplemental Figure 2. The change in averaged LFP power over the 2s baseline prior to stimulation was calculated and then grouped by genotype and sensory modality and the area under the curve of normalized power was used for statistical analysis as below.

For multi-unit spike analysis, the raw signals were band-pass–filtered (300 to 3000 Hz), after which spike detection was performed in MATLAB (R2019a). The threshold for spike detection was set to 3.5-fold of the SD of a two-second spike-free window of each recorded signal. As a quality control of the isolated multi-units, we inspected the shape of spike waveforms, and only the units with a clear negative deflection in the spike waveform were extracted. The probability distributions of the spike times around the stimulation (2s before to 3s after the stimulation) with a binning of 0.1 second were calculated and averaged for each experiment (10-20 trials per animal), and are represented as multi-unit probability (MU Probability).

### Voltage Sensitive Dye (VSD) Imaging

#### Surgery

Six month-old YAC128 Line 53 on an FVB background and wild-type FVB mice underwent a craniotomy. Mice were anesthetized with isoflurane at 3-5% for induction and maintained at 1.0-1.5% during imaging. Mice were placed on a metal plate that could be mounted on the stage of an upright microscope and the skull was fastened to a steel plate. A 7×6 mm unilateral craniotomy (bregma 2.5 to −4.5 mm, lateral 0 to 6 mm) was made and underlying dura removed as described previously^26,27^. Body temperature was maintained at 37°C with a heating pad and feedback thermistor.

#### VSD Imaging

VSD imaging was performed as described previously^25,26^. Briefly, the dye RH1692 (Optical Imaging, New York, NY) was dissolved in HEPES-buffered saline solution (1 mg ml^−1^) and applied to the exposed cortex for 60-90 min for each mouse, staining neocortical layers. VSD imaging began ~30 minutes following washing unbound VSD. The brain was covered with 1.5% agarose made in HEPES-buffered saline to minimize movement artifacts, and sealed with a glass coverslip. 12-bit images were captured at 150 Hz with a CCD camera (1M60 Pantera, Dalsa, Waterloo, ON) and EPIX E4DB frame grabber with XCAP 3.8 imaging (EPIX, Inc., Buffalo Grove, IL). The VSD was excited using a red LED (Luxeon K2, 627 nm) with excitation filters 630 +/− 15 nm. Images were recorded through a macroscope composed of front-to-front video lenses (8.6 × 8.6 mm field of view, 67μm pixel^−1^) with 1 mm depth of field. Fluorescence was filtered using a 673-703 nm bandpass optical filter (Semrock, New York, NY). For limb stimulation experiments, 10-45 trials of stimulus presentation were averaged to reduce the effects of on-going spontaneous activity. There was a 10 s interval between stimulation trials, and non-stimulation trials were collected and used for normalization of stimulated data.

#### Analysis

All VSD responses were expressed as percentage relative to baseline VSD responses ((F-F_0_)/F_0_)*100, where F_0_ is the baseline at the start of the trial, to reduce regional biases in VSD signal caused by uneven loading of the dye (calculated using MATLAB).

For region-based analyses, the coordinates of the hindlimb primary sensory area were determined by centering a 5×5 pixel ROI over the initial point of response to contralateral hindlimb stimulation for each animal. Similarly, forelimb sensory areas and barrel cortex sensory areas were experimentally mapped. The coordinates for other brain areas of interest were determined based on relative position to the hindlimb primary sensory area and stereotaxic coordinates as described previously^25,26^.

Sensory stimulations typically initiated waves of VSDI-measured activity which spread from modality appropriate sensory areas across the cortical surface. We first employed a threshold approach in Fiji-ImageJ to compare the extent of this sensory-mediated activity spread between genotypes. To do so, the exposed cortical surface in each ΔF/F VSDI movie was manually traced and the whole cortex area stored as a region of interest (ROI). The root mean square (RMS) noise of ΔF/F magnitude at each pixel within the ROI was measured during the initial 200 ms (baseline) of a movie prior to sensory stimulation. The peak ΔF/F value following sensory stimulation was identified at each pixel with a maximum signal projection of the entire movie and this value divided by a given pixel’s baseline RMS noise. The cortical area activated following sensory stimulations was determined by pixels active over a 5x baseline RMS noise threshold and compared between animals (Similar genotype differences were seen when this threshold was varied between 3 – 10x RMS noise).

Notably, the cortical area activated following sensory stimulation increased with stimulation intensity, but in the cases of forelimb and hindlimb stimulations typically plateaued at 0.5 – 1.0 mA intensities. This maximum (plateau) value was specifically examined to facilitate meaningful comparisons between animals and genotypes. Although the above analysis proved useful for quantifying the spatial extent of evoked signal spread, it could be biased by differences in baseline RMS noise between animals and only examined each pixel’s maximum ΔF/F value following stimulation. Therefore we also examined, and compared between genotypes, spatially averaged ΔF/F time-courses at select 5 × 5 pixel ROIs (outlined above) during plateau amplitude sensory stimulations.

#### Optical Flow Analysis

Optical Flow analysis of the VSD data was performed using MATLAB and Graphpad Prism. The Optical-Flow Analysis Toolbox (OFAMM) for MATLAB (http://lethbridgebraindynamics.com/ofamm/) was used to analyze the spatiotemporal dynamics of the VSD data^27^. This toolbox allows us to estimate the spatiotemporal dynamics, such as trajectory and speed, of pixels in our VSD recordings, using a variety of optical flow analysis methods. We used the Horn-Schuck (HS) method^27,61^ for our optical flow analysis, and repeated select analyses with the Combined Local-Global (CLG) method, which combines the HS method with the Lucas-Kanade (LK) method^27,62,63^. The main difference between these methods is that the LK method assumes a pixel’s motion is constant relative to neighboring pixels, whereas the HS method does not make this assumption^27^.

Optical flow analysis of 9×9 pixel (603×603 μm) regions of interest for hindlimb, forelimb, and barrel cortex primary and secondary sensory areas, as well as motor barrel cortex, was performed, to determine the trajectory and temporal speed of activity originating or flowing through that region in response to sensory stimulation. Regions were determined functionally through sensory stimulation or in relation to bregma.

#### Whole-cell voltage clamp in acute cortical slice experiments

Mice were anesthetized with isoflurane, decapitated and the brain rapidly removed. Sagittal brain slices (300μm) with sensory cortex were cut on a vibratome (VT1000 Leica) in ice cold artificial cerebrospinal fluid (aCSF, as below except 0.5mM CaCl2 and 2.5mM MgCl2) containing Kynurenic acid (1 mM) and bubbled with 95%O2/5%CO2. Slices were transferred to a holding chamber with aCSF (in mM: 125 NaCl, 2.5 KCl, 1.25 NaH2PO4, 2 CaCl2, 1 MgCl2, 25 NaHCO3, 10 glucose with osmolarity 310 mOsm) at 37°C for 40 min then kept at room temperature. Layer 2/3 pyramidal neurons were selected based on their shape and apical dendrites. For recording evoked excitatory postsynaptic currents (EPSCs) pipettes (resistance 3-5MΩ) were filled with intracellular recording solution containing (in mM:130 CsMe, 5 CsCl, 4 NaCl, 1 MgCl2, 10 HEPES, 5 EGTA, 5 QX-314Cl, 0.5 NaGTP,10 Na-phosphocreatine, 5 MgATP at pH 7.3 and 290mOsm). Stimulation was through a glass micropipette filled with aCSF positioned > 300μm ventral to the neuron. Neurons were recorded at +30mV in aCSF with added GABA_A_ receptor antagonist 50μM Picrotoxin (PTX) and 10μM AMPA/kainate receptor antagonist 6-cyano-2,3-dihydroxy-7-nitro-quinoxaline (CNQX). A stimulation train of 10 × 0.1ms at 20Hz was delivered every 30s and recorded for >5min. The GLT1/EAAT2 blocker DL-threo-b-Benzyloxyaspartic acid (TBOA, 30μM) was bath-applied for 15 min. The recording solution for spontaneous inhibitory postsynaptic currents (IPSCs) in layer 2/3 was as above with BAPTA in the internal solution and recordings were made without any drugs in the bath and neurons were held at +10mV. Neurons in layer 5 sensory cortex were held at −70mV with TTX and CNQX in the bath and using a high Cl internal solution containing (in mM: 130 CsCl, 5 NaCl, 10 HEPES, 0.5 EGTA, 4 MgATP, 0.3 Na2GTP, 5 QX-314Cl) to record miniature IPSCs.

#### Statistics

Statistical analysis was calculated with GraphPad Prism. LFP power was compared by 2-way ANOVA of area under the curve of normalized power for 1.5 s following stimulus and Šídák’s multiple comparisons test (Figures 1, 2, 5 and 6). Unpaired, two-tailed Student’s t-test was used to compare the VSDI area activated with hindlimb stimulation (Figure 3B), layer 5 mIPSC frequency and amplitude (results text), layer 2/3 membrane properties, sIPSC frequency, and amplitude (results text and Figure 4B) and isoflurane burst suppression (Supplemental Figure 1B,C,E and F). Number of cells with NMDAR events were compared by Chi-square (Figure 4A) and TBOA effects on frequency and amplitude by Kruskal-Wallis ANOVA with Dunn’s multiple comparisons post hoc (Figure 4E). P values less than 0.05 were considered significant. In whole-cell voltage clamp experiments n indicates the number of cells with the number of animals shown in brackets. For all other experiments n= number of animals. The distributions for VSDI trajectory length and maximum temporal speed (determined by optical flow analysis) were compared using the two-sample Kolmogorov-Smirnov test (Figure 3C and Supplemental Figure 4).

## Data and Code availability

The data that support the findings of this study and the code used for the analysis are available from the corresponding author upon request.

## Acknowledgements

This work was supported by resources made available through the NeuroImaging and NeuroComputation Centre at the Djavad Mowafaghian Centre for Brain Health (RRID: SCR_019086); the UBC Vice-President Research and Innovation funding for the Dynamic Brain Circuits Cluster of Excellence; Canadian Institutes of Health Research grants FDN 143210 to LAR and FDN 143209 to THM; Canadian Open Neuroscience Platform Scholar award to EK and Michael Smith Foundation for Health Research Trainee award (RT-2020-0614) to JM.

We thank Kyle Mathewson from the University of Alberta for advice and assistance with the LFP analysis and Navvab Afrashteh for assistance with Optical flow analysis. We also thank Lily Zhang, Pumin Wang and Evan Fung for technical assistance.

## Author contributions

MDS, JM, EK, DX, MHM, AC, LAR and THM designed the experiments. MDS, JM, EK, MHM, AC, A.S-D., and DR performed the research and analyzed the data. MDS, JM, EK and LAR wrote the manuscript with input from MHM, AC, DR and THM.

## Competing Interests

The authors declare no competing interests.

## Notes

### Competing Interest Statement

The authors have declared no competing interest.

