## Supplemental Figures for "Altered cortical processing of sensory input in Huntington disease mouse models"

**This PDF file includes**: Supplementary Figures 1 to 6

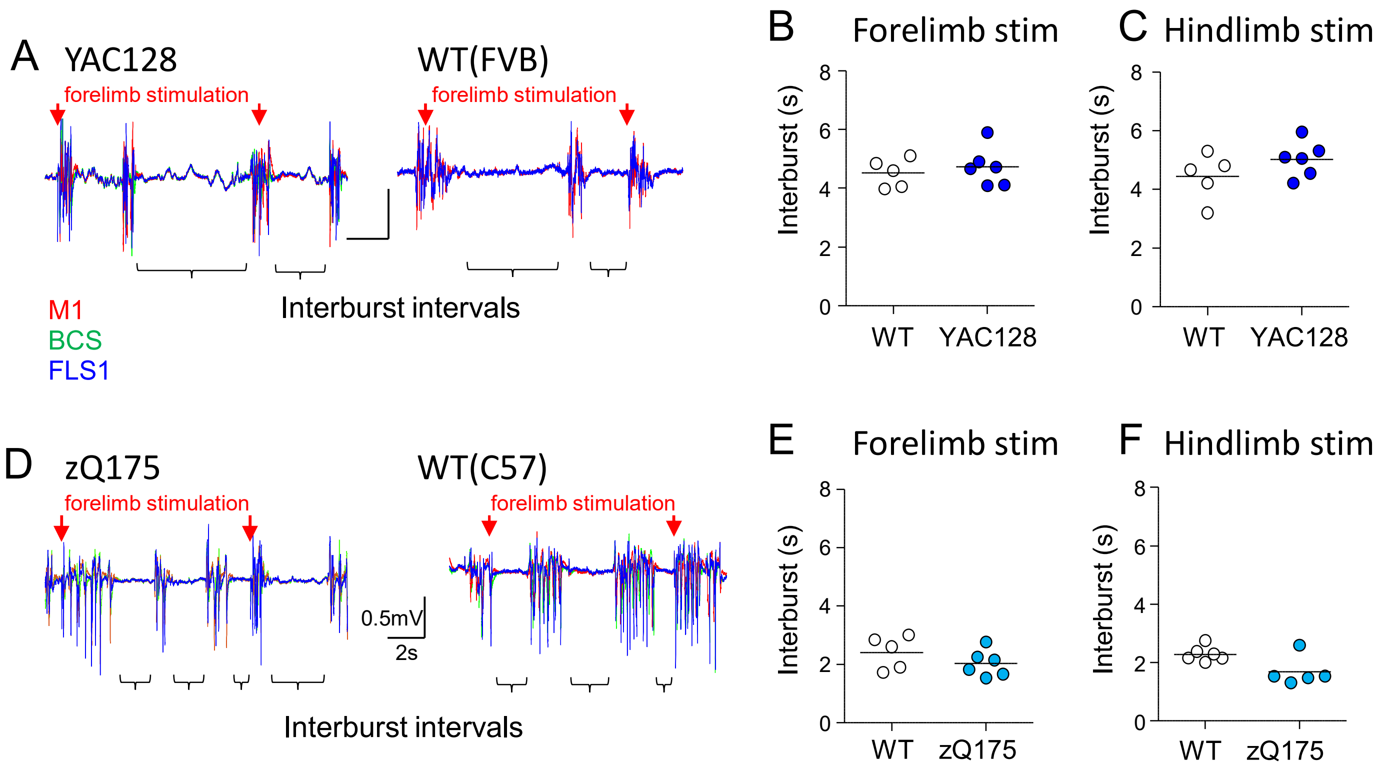


**Supplemental Figure 1.** Level of anesthetic is not significantly different between genotypes as determined by isoflurane burst suppression. A) Representative LFP recordings showing large bursts of activity following several seconds of quiet typically induced by isoflurane (interburst interval) for YAC128 (left) and WT(FVB) littermates (right). B) Genotype summary of average interburst interval (one circle per mouse) during forelimb stimulation experiments and C) hindlimb stimulation experiments. D) Representative LFP recordings showing interburst intervals for zQ715 (left) and WT(C57) littermates (right). Note lower anesthetic level with shorter intervals in zQ175 and littermates to improve detection of MU neuronal activity. E) Summary of average interburst interval during forelimb stimulation experiments and F) hindlimb stimulation experiments showing no difference between genotypes.

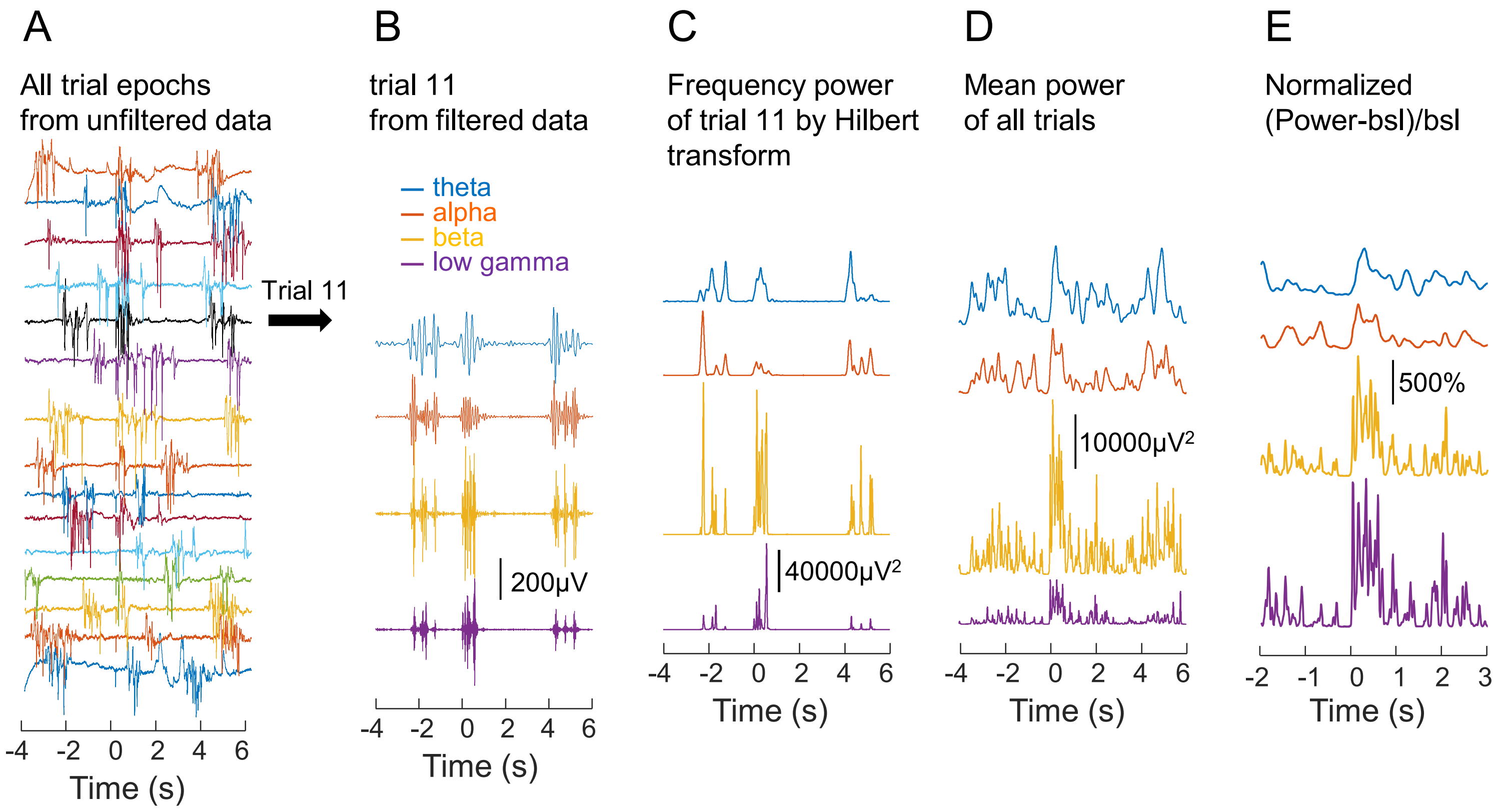


**Supplemental Figure 2.** Representative example of LFP power analysis. A) Responses to forelimb stimulation at time 0 in FLS1 showing all trial epochs from unfiltered data for comparison. B) one trial extracted after band-pass filtering the whole 5min recording. C) Hilbert transform of the same 10s trial as in B. D) the mean power of all trials for that experiment. E) Results in D normalized as the change in power divided by a 2s baseline before stimulation and 5s exported for genotype comparison.

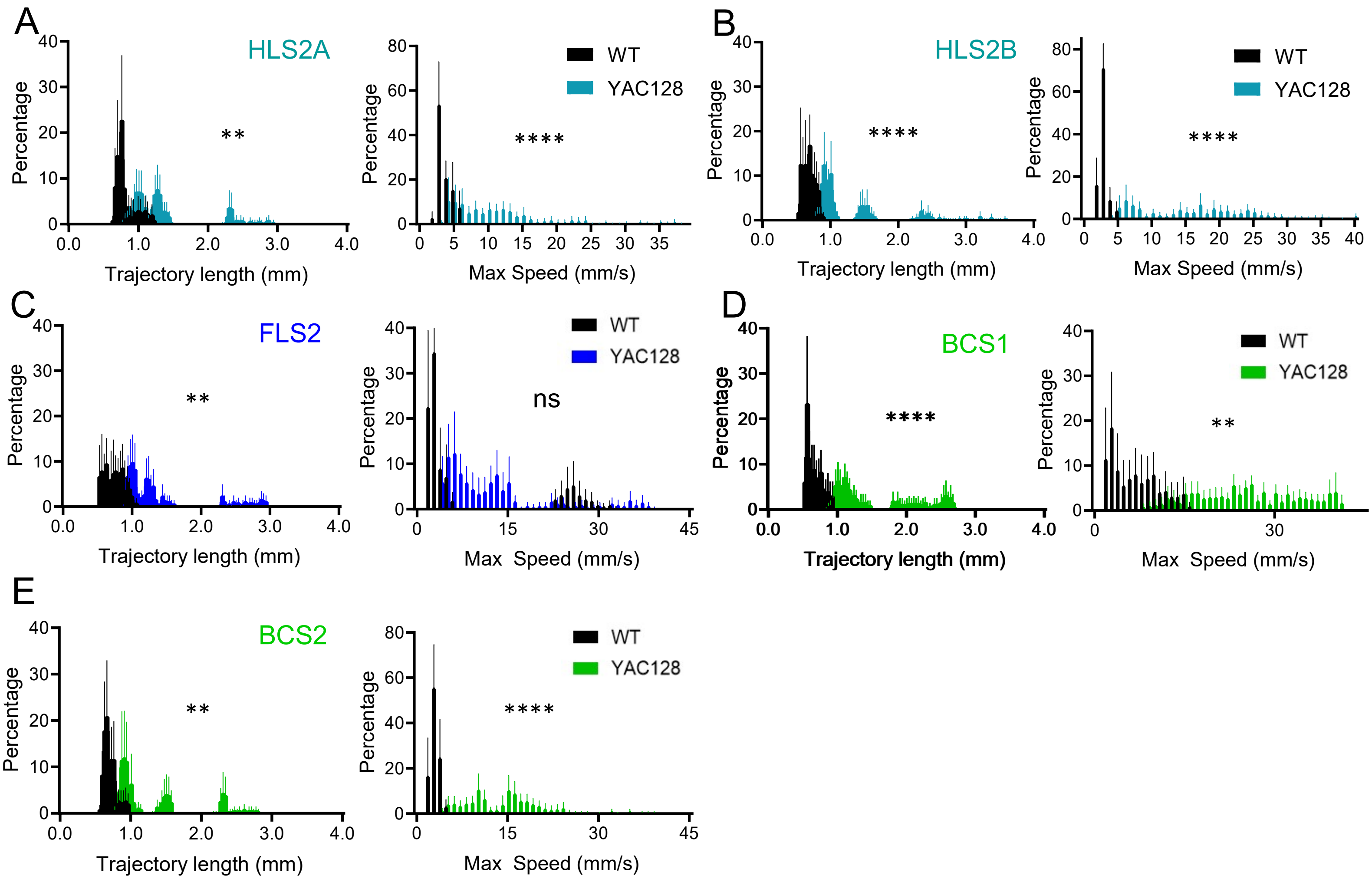


**Supplemental Figure 3.** Voltage sensitive dye imaging and optical flow analysis of functionally defined primary and secondary areas in response to hindlimb stimulation reveals increased trajectory length and maximum speed. A) Histograms showing trajectory lengths and maximum speeds reached by pixels originating in secondary hindlimb sensory area A. B) Trajectory lengths and maximum speeds of pixels originating in secondary hindlimb sensory area B. C) Trajectory lengths and maximum speeds of pixels originating in forelimb secondary sensory area. D) Trajectory lengths and maximum speeds of pixels originating in primary barrel cortex. E) Trajectory lengths and maximum speeds of pixels originating in secondary barrel cortex. *p<0.05, **p<0.01, ****p<0.0001 determined by two-sample Kolmogorov-Smirnov test.

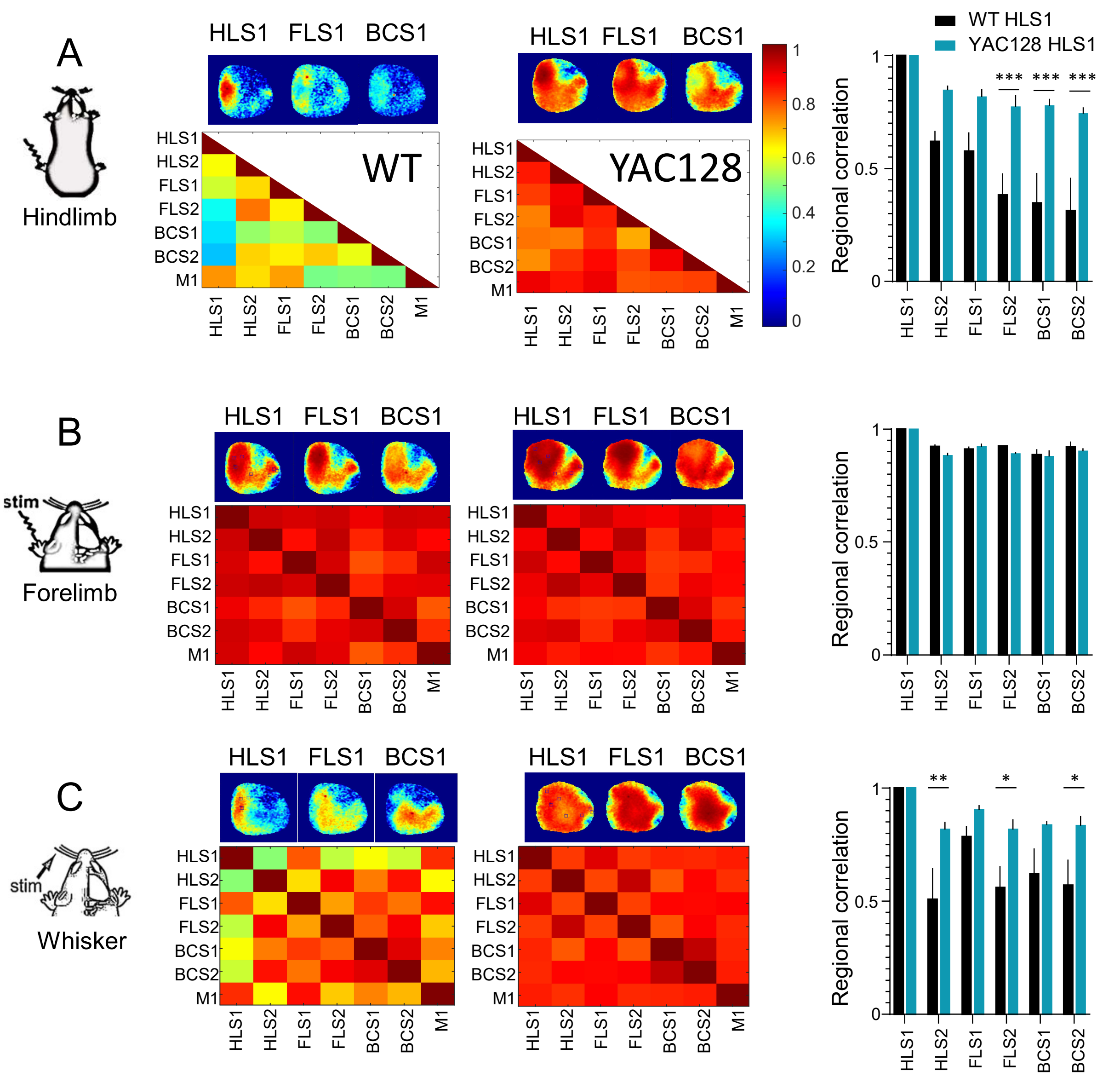


**Supplemental Figure 4.** Regional correlation of VSD activity between cortical areas is higher in YAC128 than WT following sensory stimulation. A) Representative correlation maps after hindlimb stimulation generated by a seed pixel centered in HLS1, FLS1 and BCS in WT (left) and YAC128 (right). Regional correlation matrices showing hindlimb stimulation evoked high correlation between cortical areas in YAC128 (n=4) compared to WT (n=4). HLS1 activity is significantly more correlated in YAC128 compared to WT. B) Regional correlation and representative seed-pixel-based correlation maps of VSD activity evoked by forelimb stimulation showing extensive correlation of cortical areas in both WT (left) and YAC128 (right). C) Regional correlations matrices and representative maps of cortical activity evoked by whisker stimulation in WT and YAC128. HLS1 activity is significantly more correlated in YAC128 compared to WT. n = number of mice, results are shown as mean +/- s.e.m. and ***p<0.001, **p<0.01, *p<0.05 by 2way ANOVA with Šídák's multiple comparisons test.

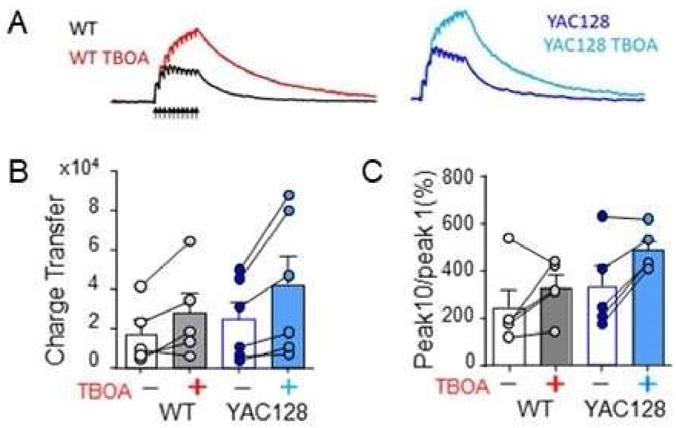


**Supplemental Figure 5.** Evoked excitatory post synaptic currents (eEPSCs) in acute brain slices with layer 2/3 sensory cortex are not significantly different between YAC128 and WT. A) Representative traces of eEPSC in response to 10 stimulations at 20Hz (arrows) in WT and YAC128 neurons before and after TBOA application to induce glutamate spillover. B) Area (charge transfer pA $ms^{-1}$) of eEPSCs after the tenth stimulation. Circles indicate change with TBOA in individual neurons. C) The ratio of the tenth peak to the first to show increase after TBOA in both WT and YAC128 neurons.

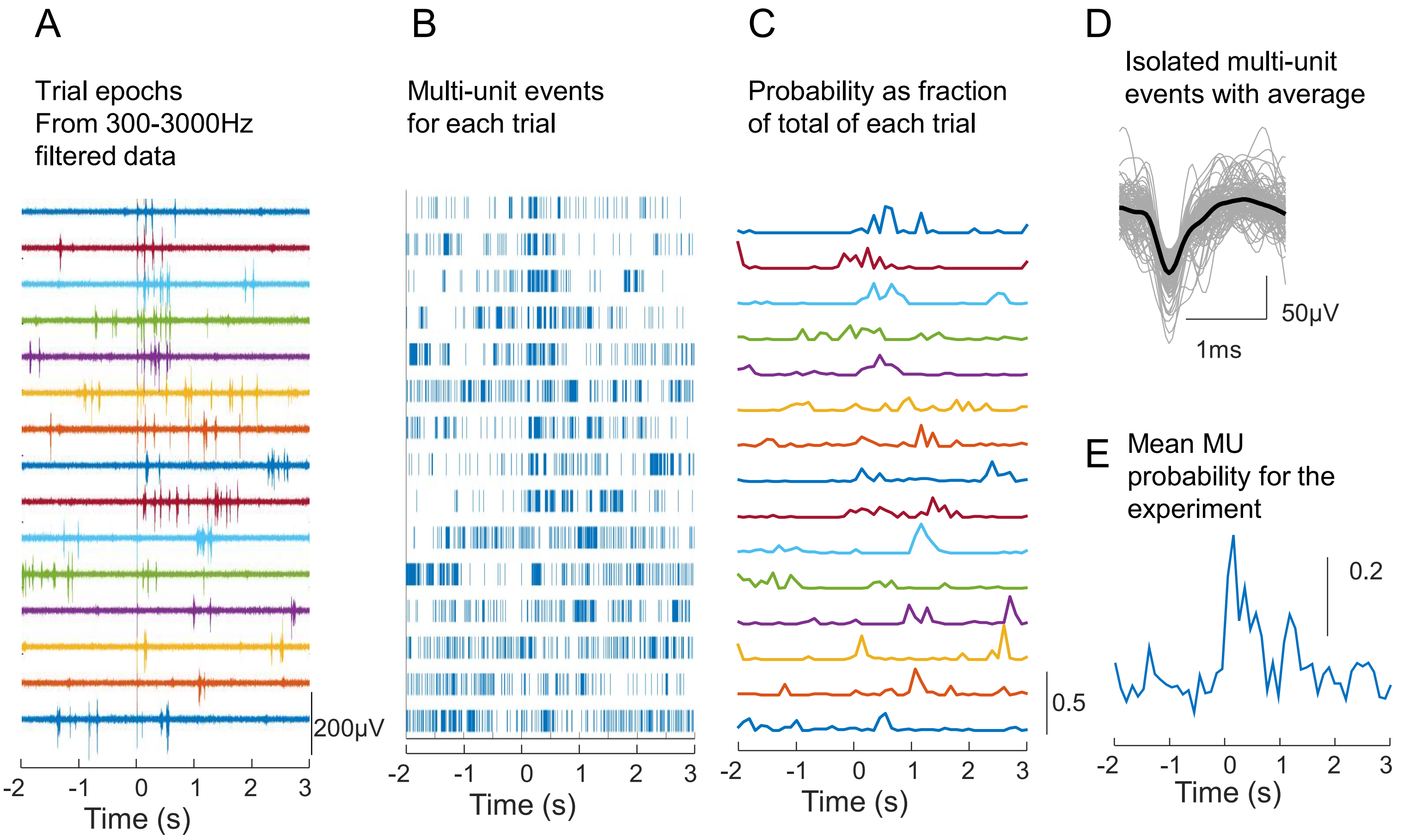


**Supplemental Figure 6.** Representative example of Multi-unit probability analysis. A) Responses to forelimb stimulation at time 0 in FLS1 showing trial epochs from filtered data. B) Spike times of all events that passed a 3.5SD baseline threshold and were interpreted as Multi-units (no sorting algorithm used). C) The probability of MU spikes in 100ms intervals as a fraction of all events in that trial. D) Isolated MU waveforms (gray) with mean waveform (black). E) Mean MU probability of all trials exported for genotype comparison.
